# Functional peroxisomes are required for heat shock-induced hormesis in *Caenorhabditis elegans*

**DOI:** 10.1101/2021.07.13.452156

**Authors:** Marina Musa, Ira Milosevic, Nuno Raimundo, Anita Krisko

**Author notes:** **Correspondence to**: Anita Krisko, Department of Experimental Neurodegeneration, University Medical Center Göttingen, Waldweg 33, 37073 Göttingen, Germany.

## Abstract

Exact mechanisms of heat shock induced lifespan extension, while documented across species, are still not well understood. Here we put forth evidence that fully functional peroxisomes are required for the activation of the canonical heat shock response and heat-induced hormesis in *C. elegans*. While during heat shock the HSP-70 chaperone is strongly upregulated in the wild-type (WT) as well as in the absence of peroxisomal catalase (*Δctl-2* mutant), the small heat shock proteins display modestly increased expression in the mutant. Nuclear localization of HSF-1 is reduced in the *Δctl-2* mutant. In addition, heat-induced lifespan extension, observed in the WT, is absent in the *Δctl-2* mutant. Activation of the antioxidant response, the pentose phosphate pathway and increased triglyceride content are the most prominent changes observed during heat shock in the WT worm, but not in the *Δctl-2* mutant. Involvement of peroxisomes in the cell-wide response to transient heat shock reported here gives new insight into the role of organelle communication in the organisms stress response.

## Introduction

Heat shock response (HSR) is the cellular response to stress, characterized by robust upregulation of heat shock proteins or chaperones under control of transcription factor HSF-1 in *C. elegans*. It is necessary for maintenance of proteostasis, disruption of which is associated with several human diseases, including Parkinson’s, Alzheimer’s, and even some forms of cancer (Morimoto 2008, Mendillo, Santagata et al. 2012). The adaptive stress response in recent years has been studied in the context of aging and disease, and many hormetic treatments have been demonstrated to be beneficial for survival. These beneficial effects have generally been attributed to the activation stress response pathways, which depend on the type of the stressor and downstream signaling targets (Cypser and Johnson 2002, Calabrese, Bachmann et al. 2007, Rattan and Demirovic 2009).

Transient exposure to heat, irradiation and oxidative stress have all been shown to be able to extend lifespan across organisms from yeast to rodents (Shama, Lai et al. 1998, Olsen, Vantipalli et al. 2006, López-Martínez and Hahn 2014, Scott 2014, Baldi, Bolognesi et al. 2017, Cox, McKay et al. 2018, Musa, Perić et al. 2018). Although the connection between stress and hormesis is well described, the exact mechanisms are still somewhat of a debate. Canonically, hormetic effects are attributed to the increased concentration of chaperones inside the cell, but there is mounting evidence that tells a more complex story. Several pathways aside from HSR are needed for the hormetic effect to occur, including inhibition of TORC1 and activation of autophagy, increase in respiration, and accumulation of mitochondrial ROS (Olsen, Vantipalli et al. 2006, Bonawitz, Chatenay-Lapointe et al. 2007, Pan and Shadel 2009, Van Raamsdonk and Hekimi 2009, Yang and Hekimi 2010, Kumsta, Chang et al. 2017, Perić, Lovrić et al. 2017, Musa, Perić et al. 2018).

The bulk of what we know about the systemic stress response and hormesis comes from studies done on budding yeast, which remains an invaluable model system for studying the fundamental mechanisms of regulation of stress responses. However, stress responses in yeast are cell autonomous and relatively simple when compared to multicellular organisms such as *C. elegans*, where stress responses are cell nonautonomous and need to be regulated in a tissue specific manner through a series of negative and positive regulators (Prahlad, Cornelius et al. 2008, Guisbert, Czyz et al. 2013, van Oosten-Hawle, Porter et al. 2013, van Oosten-Hawle and Morimoto 2014, Takeuchi, Suzuki et al. 2015). In *C. elegans*, increases in temperature are sensed by the thermosensory AFD neurons. The exact mechanisms of how these neurons translate the temperature change signals to other cells and distal tissues is yet to be fully described.

Peroxisomes, oxidative organelles that play a role in oxidation of very long fatty acids and, in general, cell metabolism, have a number of vital roles that may affect survival during stressful conditions, including but not limited to their connection with mitochondria, already implicated in many aspects of stress biology. Peroxisomal catalase *ctl-2* mutant displays a shortened lifespan (Petriv and Rachubinski 2004), and peroxisomes are known to change in shape, size and abundance in response to environmental stimuli and cell status.

Here, we present results revealing that peroxisomes play a crucial role for the HS-induced lifespan extension in *C. elegans*. Moreover, our results suggest functional peroxisomes are necessary for proper heat shock response activation. While, traditionally, the role of heat shock proteins has been put forward as most relevant for heat-induced hormesis, our results reveal an unexpected alliance between the heat shock response and peroxisomes, hubs of ROS and fatty acid metabolism.

## Results

### Peroxisomal mutants have impaired HSR and thermotolerance

We aimed to investigate the metabolic response of *C. elegans* to mild HS and a putative role for peroxisomes in this process. We focused on two *C. elegans* strains: WT and peroxisomal mutant *Δctl-2*. The *Δctl-2* mutant, which lacks the peroxisomal catalase *ctl-2*, was chosen because it displays a shortened lifespan (Petriv and Rachubinski 2004). Peroxisomal catalase *ctl-2* is the only peroxisomal catalase and is responsible for about 80% of all catalase activity in *C. elegans* (Petriv and Rachubinski 2004). The 4-hour HS at 30°C was administered in L4 stage while control worms were kept at 20°C throughout the experiment. In WT worms, we measured increased median and maximum lifespan; median lifespan was increased from 11 to 14 days post HS, and maximum lifespan from 17 to 20 days post HS (Figure 1A). In contrast, in *Δctl-2* we measured a median lifespan of 10 days in both control and HS group, while maximum lifespan was 13 and 15 days post HS for control and HS group respectively (Figure 1B). Therefore, mild HS had no effect on the *Δctl-2* strain lifespan. Concurrent with other reports of increased lifespan at the cost of fitness (Van Raamsdonk and Hekimi 2009, Lapierre, Gelino et al. 2011, Hansen, Flatt et al. 2013), increased lifespan following HS treatment in WT was accompanied by modestly smaller brood size. The *Δctl-2* worms displayed a strong decline in brood size compared to the WT, as well as a further decline due to HS, despite not exhibiting an extended lifespan (Figure 1C).

**Figure 1.**
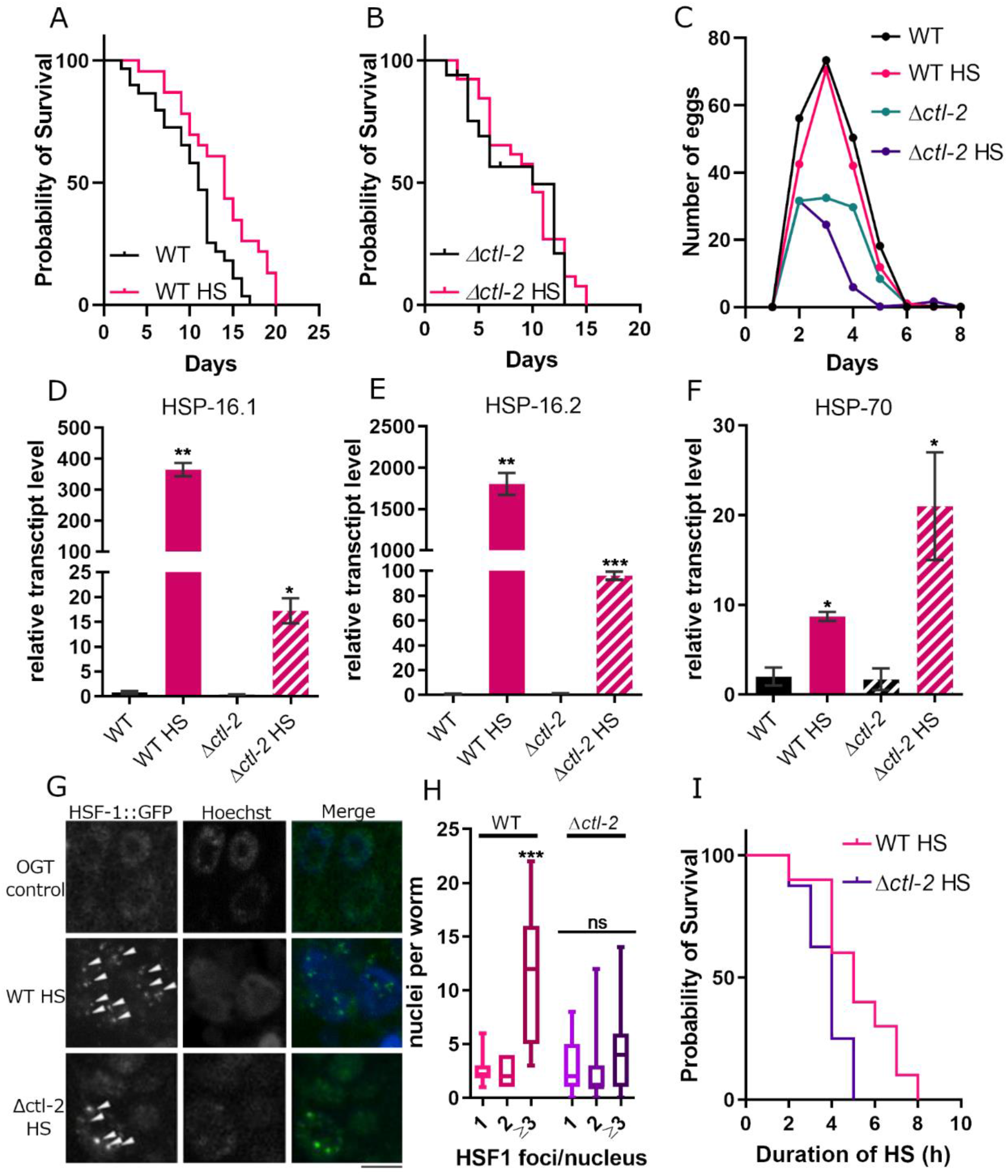
Heat shock response is constrained in peroxisomal *Δctl-2* mutant. (A) Mild transient HS at L4 stage extends WT lifespan. Median lifespan following HS is 14 days compared to 11 days at OGT, and maximum lifespan is extended from 17 days to 20 days after HS. (B) In *Δctl-2* background lifespan is not affected by HS; median lifespan is 10 days for both conditions, and maximum lifespan is 13 days at OGT and 15 days following HS treatment. HS is administered at day 0. P value is >0.05 (Mantel-Cox). (C) *Δctl-2* worms has reduced brood size compared to WT. (D-F) HSR is activated in both strains after 4-hour HS, however, transcript levels of *hsp16*.*1* and *hsp16*.*2* are lower in *Δctl-2* compared to WT. HSP-70 expression is higher in *Δctl-2* mutant compared to WT following HS. Gene expression normalized to *act-1*. Error bars are mean ± SD. \*\*\*\**P*<0.0001; \*\*\**P* < 0.001; \*\**P* < 0.01; \**P* < 0.05 (t-test). (G) HSF-1::GFP forms foci in the nucleus during HS in the WT. The black bar represents 5 μm. (H) Quantification of images displayed in (G) shows that *Δctl-2* mutants have a significantly lower proportion of nuclei with more than 3 foci: WT averages 15 nuclei with 3 or more HSF1 puncta after 4 hours of HS, compared to only 4 nuclei in *Δctl-2*. \*\*\**P* < 0.001; \*\**P* < 0.01; \**P* < 0.05 (ANOVA). (I) *Δctl-2* mutant shows decreased ability to withstand heat stress. \*\*\**P* < 0.001; \*\**P* < 0.01; \**P* < 0.05 (Mantel-Cox).

As robust upregulation of the expression of chaperones/heat shock proteins (HSP) is the major hallmark of HSR, we measured transcription levels of several HSPs. Surprisingly, we see that while small HSPs (sHSPs) were robustly upregulated in WT during mild HS, their upregulation was significantly less pronounced in the *Δctl-2* mutant (Figure 1D,E). Interestingly, the expression of HSP16.2 in young worms in response to stress has been positively correlated with lifespan (Rea, Wu et al. 2005). On the other hand, transcript levels of the HSP-70 protein were almost twice as abundant in the *Δctl-2* mutant compared to WT (Figure 1F), further suggesting that the transcriptional response to HS in *Δctl-2* mutants is aberrant.

The upregulation of HSPs during HS is coordinated by the transcription factor HSF-1, which is usually kept inactive by binding to chaperones. When the levels of unfolded proteins increase, the chaperones are recruited to these proteins, releasing HSF-1, which forms a homotrimer that binds DNA in sequences known as heat shock elements (HSEs), from where it stimulates transcription. The binding of HSF-1 to the HSEs can be visualized by a HSF-1::GFP fusion protein; the HSF-1::GFP bound to sites of HSEs is visible as GFP foci in the nucleus, as opposed to diffuse green signal of unbound protein (Morton and Lamitina 2013). Quantification of the HSF-1::GFP foci showed fewer foci in the *Δctl-2* worms compared to WT (Figure 1G,H), suggesting altered regulation of the HSR in the peroxisomal mutant. The decreased activation of HSR and lower upregulation of HSPs is also supported by the decreased thermotolerance of the *Δctl-2* strain: survival measurement at the restrictive temperature of 37°C showed that WT worms are more heat tolerant compared to *Δctl-2* (Figure 1I).

### Aberrant stress responses in peroxisomal mutants

HS causes a range of changes inside the cell, from misfolding proteins to metabolic changes. Therefore, we explored in more detail the expression levels of target genes from both aspects. We found that the unfolded protein response (UPR) was active during HS in WT but was relatively modest in the *Δctl-2* mutant (Figure 2A). While in WT the expression of the UPR markers, UGGT-1, PDI-6 and CRT-1, is strongly increased, in the *Δctl-2* mutant the increase is less prominent. Mitochondrial UPR (mt UPR) on the other hand was not activated in *Δctl-2* strain during HS, while in WT it was evidenced solely by upregulation peptidase CLPP-1 (Figure 2B). It was reported previously by Ben Zvi and colleagues (Ben-Zvi, Miller et al. 2009) that HSR and UPR activation is constrained in older animals, which would support the idea that *Δctl-2* strain exhibits a progeric-like phenotype (Petriv and Rachubinski 2004).

**Figure 2.**
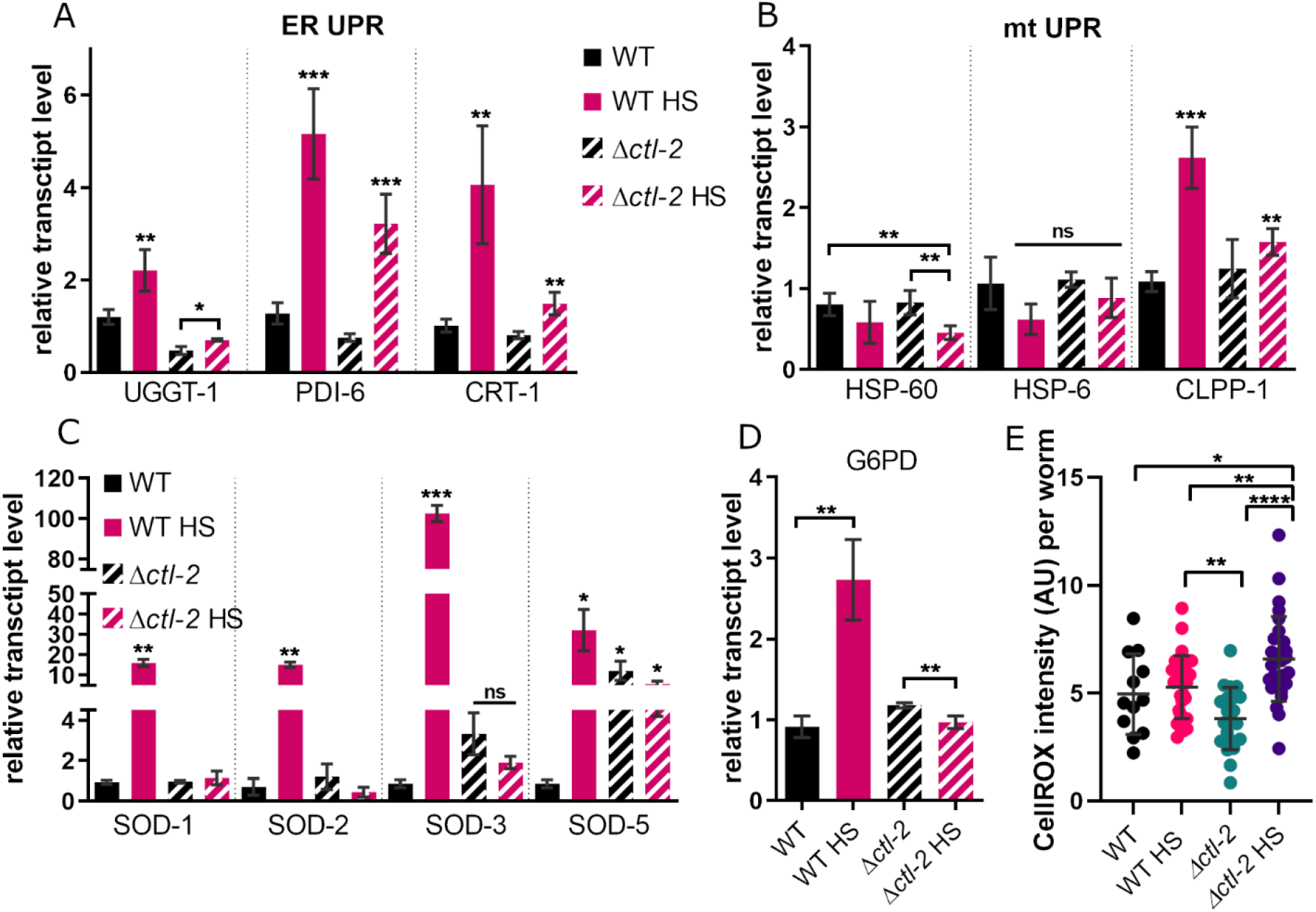
HS elicits distinct metabolic changes in WT and *Δctl-2* strain. (A) ER UPR was activated in both WT and *Δctl-2* strains. (B) CLPP-1 caspase of the mtUPR was modestly upregulated in WT, but not in *Δctl-2* strain. (C) HS does not affect transcript levels of SOD enzymes in the *Δctl-2* strain, but leads to strong upregulation in the WT. (D) HS triggers an increase in transcript levels of G6PD, rate limiting enzyme of the pentose phosphate pathway, in WT but not in *Δctl-2* strain. Gene expression normalized to ACT-1. Error bars are mean ± SD. (E) ROS levels are comparable between WT and *Δctl-2* mutant at OGT but are increased in *Δctl-2* during HS. ****P<0.0001; ***P < 0.001; **P < 0.01; *P < 0.05 (ANOVA).

We have previously shown that yeast experience oxidative stress during HS (Musa, Perić et al. 2018). This seems to be the case also in WT *C. elegans*, as suggested by multiple parameters measured, including the increase in transcript levels of SOD enzymes (Figure 2C). RT-qPCR measurement of superoxide dismutase (SOD) enzymes revealed that both cytosolic (SOD-1 and SOD-5) and mitochondrial (SOD-2 and SOD-3) superoxide dismutases were upregulated in the WT during HS, while HS did not affect transcript levels of these enzymes in the *Δctl-2* strain. However, transcript levels of the cytosolic SOD-5 in *Δctl-2* strain were increased already at OGT compared to WT and were comparable to WT HS levels (Figure 2C).

Pentose phosphate pathway or PPP is generally associated with oxidative stress, but can also be induced during heat stress as a consequence of increased superoxide production (Stincone, Prigione et al. 2015, Musa, Perić et al. 2018). While transcript levels of PPP rate limiting enzyme G6PD-1 as measured by RT-qPCR were expectedly increased in WT during HS (Stincone, Prigione et al. 2015, Musa, Perić et al. 2018), this was not the case in *Δctl-2* mutant suggesting that *Δctl-2* strain did not activate PPP during HS (Figure 2D). Lack of upregulation of key antioxidative enzymes during HS indicates that *Δctl-2* strain does not fully sense oxidative stress during HS, presumably as a consequence of catalase deficiency, or is not able to respond appropriately. As several enzymes seem constitutively upregulated in *Δctl-2* strain (Figure 2C), it is possible that lack of further upregulation of chaperones is an adaptation to avoid toxic effects of prolonged activation of stress response pathways, and that the threshold for their activation is increased.

To estimate the amount of ROS in *C. elegans*, we used CellROX dye, whose fluorescence is proportional to the amount of ROS in the cell. Interestingly, the CellROX fluorescence in WT was not affected by HS, likely as a result of the activation of antioxidant defenses. In the *Δctl-2* strain, the ROS levels were comparable to the WT strain at OGT, as indicated by CellROX fluorescence, but were significantly increased following HS (Figure 2E). Therefore, it is plausible that the high levels of SOD enzymes in the WT are efficiently containing the ROS levels and preventing their accumulation during HS, while the increasing ROS levels in the *Δctl-2* strain during HS are not neutralized. The mechanism preventing the *Δctl-2* mutant to activate the antioxidant defenses remains to be elucidated.

### Peroxisome number and morphology is affected by HS

Peroxisomes change in shape, size and abundance in response to environmental stimuli and cell status. Given the differences observed in the response to HS between the WT and the *Δctl-2* mutant, we turned our attention to the peroxisome morphology and function. In this context, the peroxisome import machinery is important for proper assembly of the organelle. We measured the expression levels of two peroxisomal transport proteins, PRX-5 and PRX-11, and found no differences in expression between WT and *Δctl-2* strains, suggesting that morphogenesis should not be impaired and that proteins needed for their proper function should be present, with the exception of *ctl-2* in the *Δctl-2* strain (Figure 3A).

**Figure 3.**
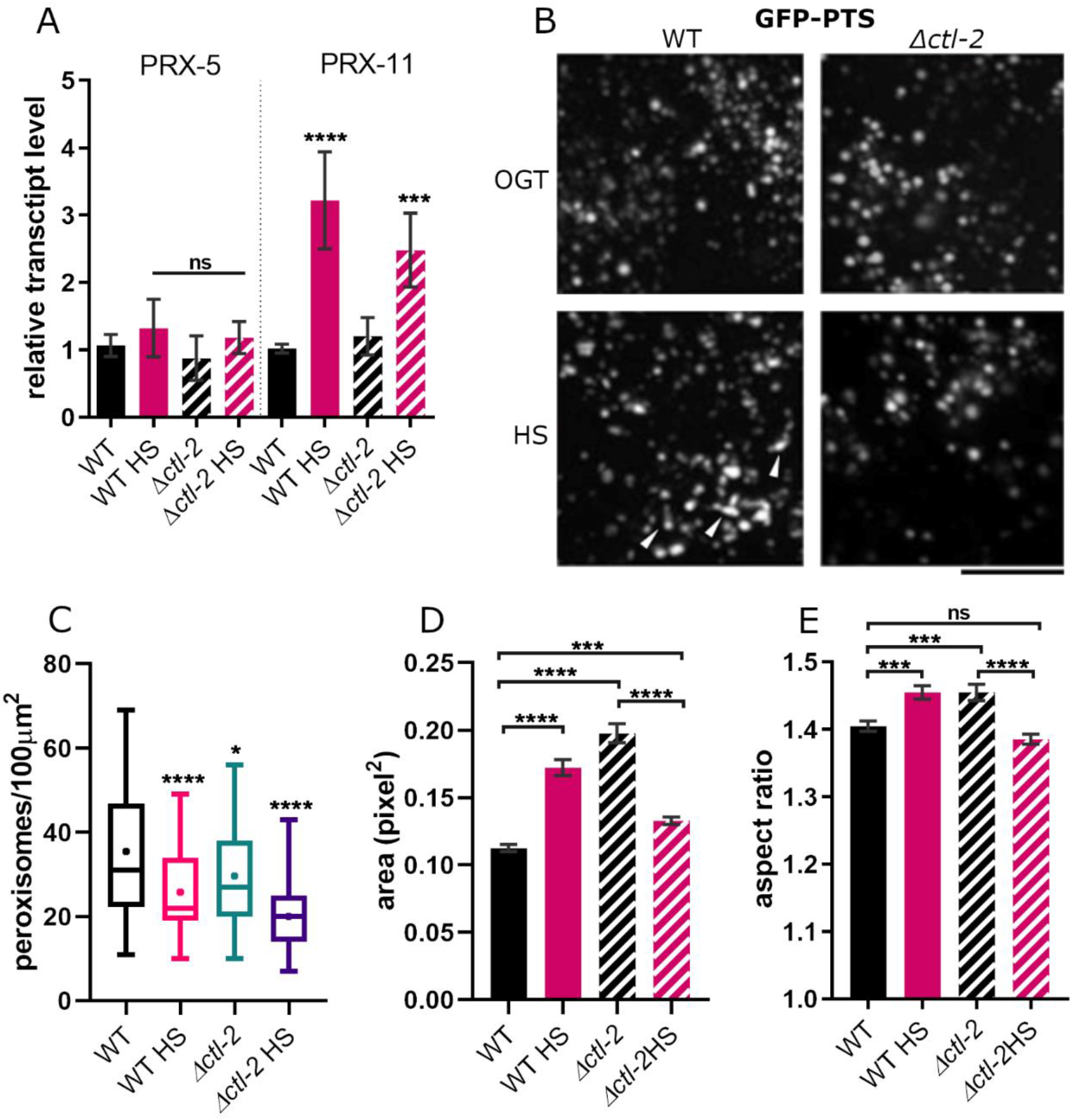
Peroxisome morphology and number are affected by mild transient HS. (A) Transcript levels of main peroxisome biogenesis proteins responsible for its content import is similar in WT and *Δctl-2*, both in control and HS-treated samples, suggesting no significant difference in peroxisomal import between the two strains. Gene expression normalized to *act-1*. Error bars are mean ± SD. \*\*\*\**P*<0.0001; \*\*\**P* < 0.001; \*\**P* < 0.01; \**P* < 0.05 (ANOVA). (B) Representative images of peroxisomes visualized using a GFP-PTS fusion protein. The black bar represents 5 μm. (C) Analysis of the images displayed in (B) shows that, on average, WT worms have more peroxisomes than *Δctl-2* strain. Their number is reduced during HS. The symbol signifies the mean. (D) *Δctl-2* peroxisomes are on average larger than those in WT; during HS WT peroxisomes increase in size while *Δctl-2* peroxisomes become smaller. (E) Aspect ratio of the analyzed peroxisomes shows that WT peroxisomes elongate during HS, while in *Δctl-2* the trend is opposite. The graphs represent mean ± SEM. ****P<0.0001; ***P < 0.001; **P < 0.01; *P < 0.05 (ANOVA).

Peroxisomes were visualized using GFP fused with peroxisomal import signal (PTS::GFP; Figure 3B). Already at OGT, in the *Δctl-2* strain peroxisomes were less abundant, or less able to import proteins, than in the WT (Figure 3C). Moreover, imaging revealed decreased numbers of peroxisomes following HS in both WT and *Δctl-2* strain compared to OGT of each strain (Figure 3C). *Δctl-2* peroxisomes were also characterized by a larger size compared to WT at OGT, measured as peroxisome area in square pixels (Figure 3D), consistent with previous reports (Petriv and Rachubinski 2004). Interestingly, while WT peroxisomes tended to increase in size during HS, the size of *Δctl-2* peroxisomes decreased (Figure 3D).

We also observed subtle changes in peroxisome shape between WT and *Δctl-2* strain. To assess changes in the peroxisome shape between the strains we measured them along the shortest and longest axis, the ratio of which approximates their sphericity: aspect ratio of 1 would describe a perfect sphere while increasing values indicate elongation. At OGT, the WT peroxisomes appeared slightly more spherical than *Δctl-2* peroxisomes with ∼4.5% increase in the aspect ratio in the *Δctl-2* mutant (Figure 3E). During HS, the sphericity of the WT peroxisomes was reduced with the aspect ratio increased by ∼5%, suggesting the elongation of the peroxisomes (Figure 3E). On the other hand, the *Δctl-2* peroxisomes displayed a tendency to become more spherical during HS (Figure 3E) with the aspect ratio decrease of ∼6% compared to the same strain at OGT. More experiments are necessary to dissect the causes and the importance of these changes because, despite being subtle, they might point towards impaired peroxisomal function in the *Δctl-2* mutant.

### Fatty acid metabolism and storage are affected by HS and Δctl-2 mutation

One of the most important processes taking place in the peroxisomes is oxidation of very long fatty acids, products of which are stored as a reserve energy supply or shuttled to mitochondria to be fully oxidized and used to produce ATP. While mitochondria can oxidize most FAs themselves, very long chain FAs can only be oxidized in the peroxisomes. For example, phytanic acid is converted to pristanic acid via α-oxidation, then broken down further through multiple rounds of β-oxidation in the peroxisome to FAs that can be fully oxidized in mitochondria. Therefore, changes in the peroxisome metabolism could cause changes in substrate availability for the mitochondrial β-oxidation. We found that peroxisomal β-oxidation is suppressed during HS in both WT and *Δctl-2*, evidenced by downregulation of the gene encoding peroxisomal straight-chain acyl-CoA oxidase, ACOX-1, the rate limiting enzyme for very long chain FA β-oxidation (Figure 4A). Following HS, both WT and *Δctl-2* strains displayed ∼25% of the transcript levels measured at OGT (Figure 4A). A similar trend was observed in the second step of peroxisomal β-oxidation, catalyzed by MAO-C-like dehydratase domain protein, MAOC-1 (Figure 4B). However, the last step, the thiolytic cleavage of 3-ketoacyl-CoA catalyzed by DAF-22 (propanoyl-CoA C-acyltransferase), was unaffected by HS in WT. Interestingly, in *Δctl-2* strain the transcript levels for enzyme DAF-22 were increased twofold at OGT compared to WT and decreased significantly following HS (Figure 4C). These results may suggest that this aspect of peroxisomal function may be important in the context of response to HS. The differences in the transcript levels of β-oxidation enzymes between WT and *Δctl-2* strain leave open a possibility that the levels of β-oxidation intermediates and final products are changed in the *Δctl-2* mutant compared to WT, thus possibly affecting both mitochondrial function and the energy supply of the cells in form of fatty acids.

**Figure 4.**
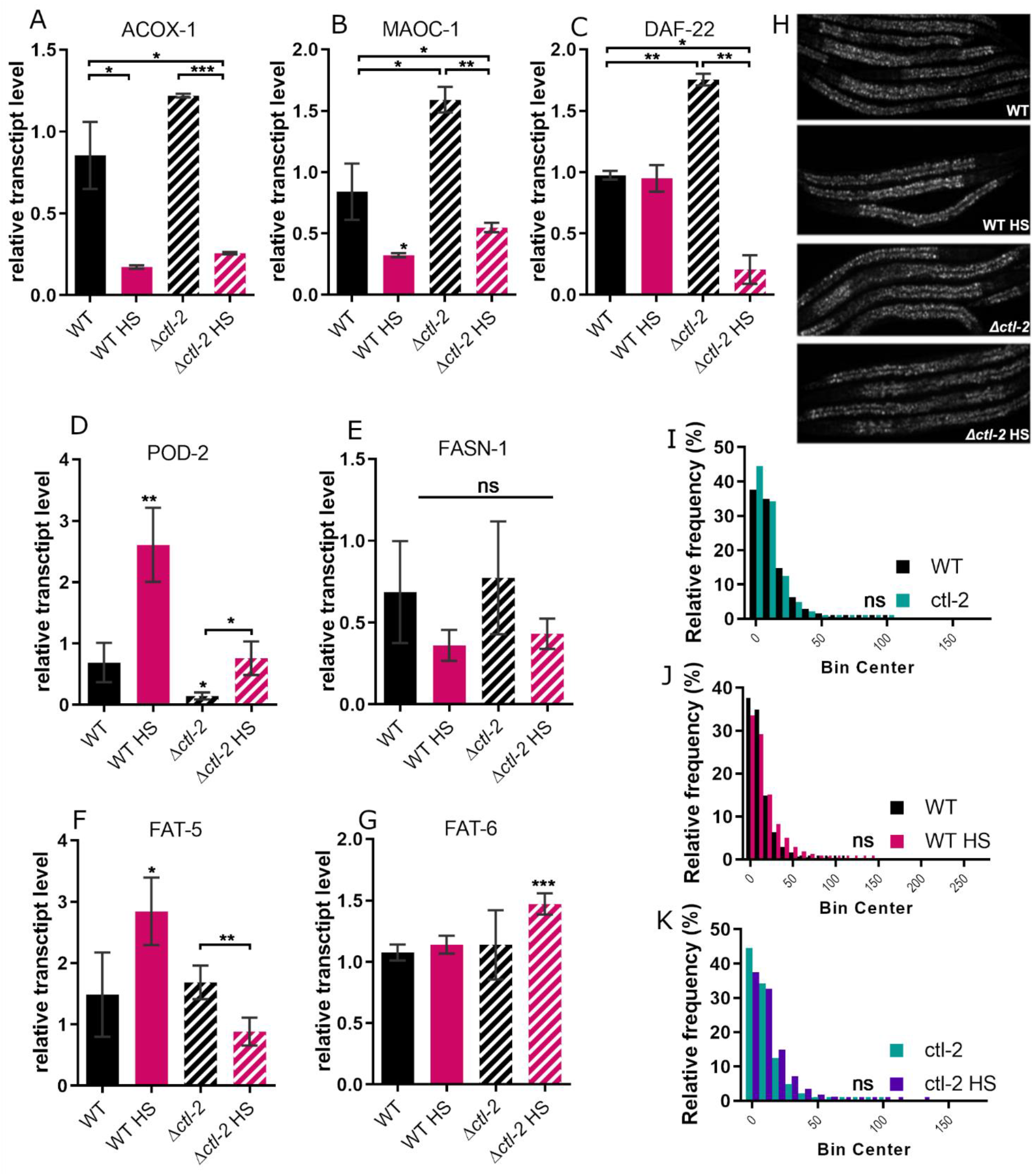
Fatty acid metabolism is affected by the absence of the peroxisomal catalase. Major peroxisomal FA β-oxidation enzymes (A) ACOX-1, (B) MAOC-1, and (C) DAF-22 expression levels are increased in the *Δctl-2* mutant at OGT compared to WT. (D-G) *Δctl-2* mutant displays different regulation of genes involved in fatty acid synthesis during HS compared to WT. Gene expression normalized to ACT-1. Error bars are mean ± SD. ****P<0.0001; ***P < 0.001; **P < 0.01; *P < 0.05 (ANOVA). (H) Representative images of Nile red staining of lipid droplets in entire worms. (I-K) The size of lipid droplets from (H) is not significantly different between WT and *Δctl-2* mutant and was not affected by HS. Bin center represents lipid droplet area measured in pixels. ****P<0.0001; ***P < 0.001; **P < 0.01; *P < 0.05 (Mann-Whitney).

Proper metabolism of FAs is essential, especially in the context of HS, as lipids are major components of the cellular membranes. The membrane alters its composition to adapt to variations in environmental conditions and deal with different stressors, such as temperature. Since our results suggest that FA β-oxidation is affected by HS, we further aimed to assess the FA synthesis and storage. To that end, we measured the transcript levels of major FA synthesis enzymes and we visualized FA stores in fixed worms. We found distinct differences in the transcript levels of the rate limiting enzyme acetyl-CoA carboxylase (POD-2), which was roughly tripled at HS in both strains, but the levels of POD-2 coding transcripts were also significantly lower at OGT in *Δctl-2* compared with WT (Figure 4D). While there were no significant changes in transcript levels of fatty acid synthase (FASN-1) between WT and *Δctl-2* (Figure 4E), Δ9 desaturases FAT-5 and FAT-6 were regulated differently. More specifically, FAT-5 expression roughly doubled following HS in WT while in the *Δctl-2* mutant it was halved (Figure 4F). Furthermore, the FAT-6 expression was unchanged following HS in WT while in the *Δctl-2* mutant was increased by ∼25% (Figure 4G). As FAT-5 and FAT-6 have different substrate specificities (FAT-5 readily desaturates palmitic acid (16:0) but not stearic acid (18:0) which is desaturated by FAT-6), the differences in their expression between strains suggest that the proportions of different FA species in the two strains could be impacted. As lipid composition is crucial, especially in conditions of changing temperature, these changes could have major impact on the worm’s survival during HS.

FAs are stored mainly in the *C. elegans* intestine inside so called lipid droplets. Lipid droplets can be visualized using Nile Red dye, which stains neutral lipids (Figure 4H). Nile Red staining revealed no significant difference between strains, nor any discernable effect of the HS treatment (Figure 4I-K). However, Nile Red does not differentiate between different lipid species. Therefore, we stained the worms using Oil Red O dye, which stains triglycerides (Figure 5A, B). The Oil Red O staining was not significantly affected by HS, but revealed a slight difference between the two strains, which suggests that triglyceride production and/or storage is affected in the *Δctl-2* worms compared to WT, as they retained less of the Oil Red O dye, indicating lesser triglyceride content in the *Δctl-2* mutant (Figure 5C-K). This is consistent with the RT-qPCR results revealing differences in FA synthesis between the two strains, as decreased transcription of POD-2 is usually associated with decreased triglyceride levels (Kim, Grant et al. 2016). While further experiments are needed to investigate the details of the lipid production, storage, and utilization in the *Δctl-2* strain, this data provides evidence that there are differences in the FA content between the peroxisomal mutant and WT.

**Figure 5.**
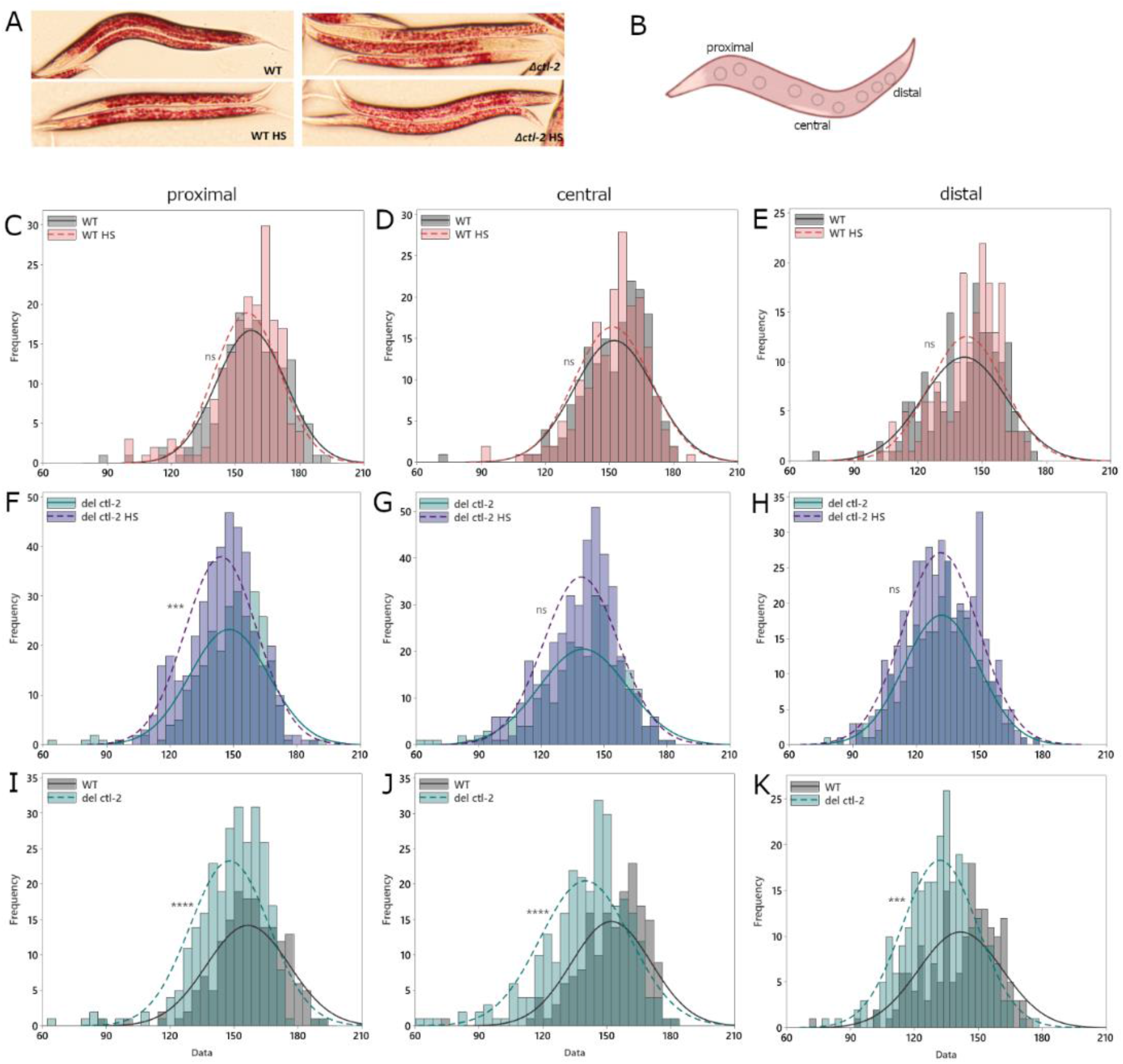
Triglyceride content is reduced in *Δctl-2* strain. (A) Representative images of Oil Red O stained worms. (B) Staining intensity is measured separately in the area between the pharynx and the vulva (proximal), area around the vulva (central), and the tail section (distal), avoiding the darker edge of the worm and the vulva, as integrated density i.e. sum of values of pixel intensities normalized to area. Three measurements of each section are averaged for each worm. (C-K) *Δctl-2* worms show lower intensity Oil Red O staining compared to WT, indicating lower triglyceride content. ****P<0.0001; ***P < 0.001; **P < 0.01; *P < 0.05 (Mann-Whitney).

### Mitochondrial morphology is altered in Δctl-2 mutant during HS

Given the metabolic connections between peroxisomes and mitochondria, and the changes we observed in peroxisomes, we turned to mitochondria next. Mitochondria form physical contact sites with peroxisomes in order to communicate and facilitate exchange of molecules such as lipids from peroxisomes that are to be used by mitochondria to produce ATP. Mitochondria were visualized with MitoTracker Deep Red dye and imaged in live worms using spinning disc confocal microscope (Figure 6A). Analysis of stained mitochondria indicated that WT and *Δctl-2* mitochondria displayed differences in morphology based on the branching of imaged mitochondria. The number of branches, branch junctions, and the average branch length were used to describe mitochondrial networks, while individual mitochondria were separated based on their shape. The analysis of the mitochondrial networks shows that the average *Δctl-2* strain mitochondrion is less branched, with fewer, shorter branches compared with WT. WT mitochondrial branches seem to be ∼25% longer than those observed in *Δctl-2* strain at OGT. Following HS, however, the average branch length is decreased by ∼50% in WT, while it is increased by ∼20% in *Δctl-2* strain (Figure 6B-D). In both WT and *Δctl-2* strains mitochondria appear less branched following HS compared to OGT, indicated by the decrease in number of branches and branch junctions (Figure 6E-J).

**Figure 6.**
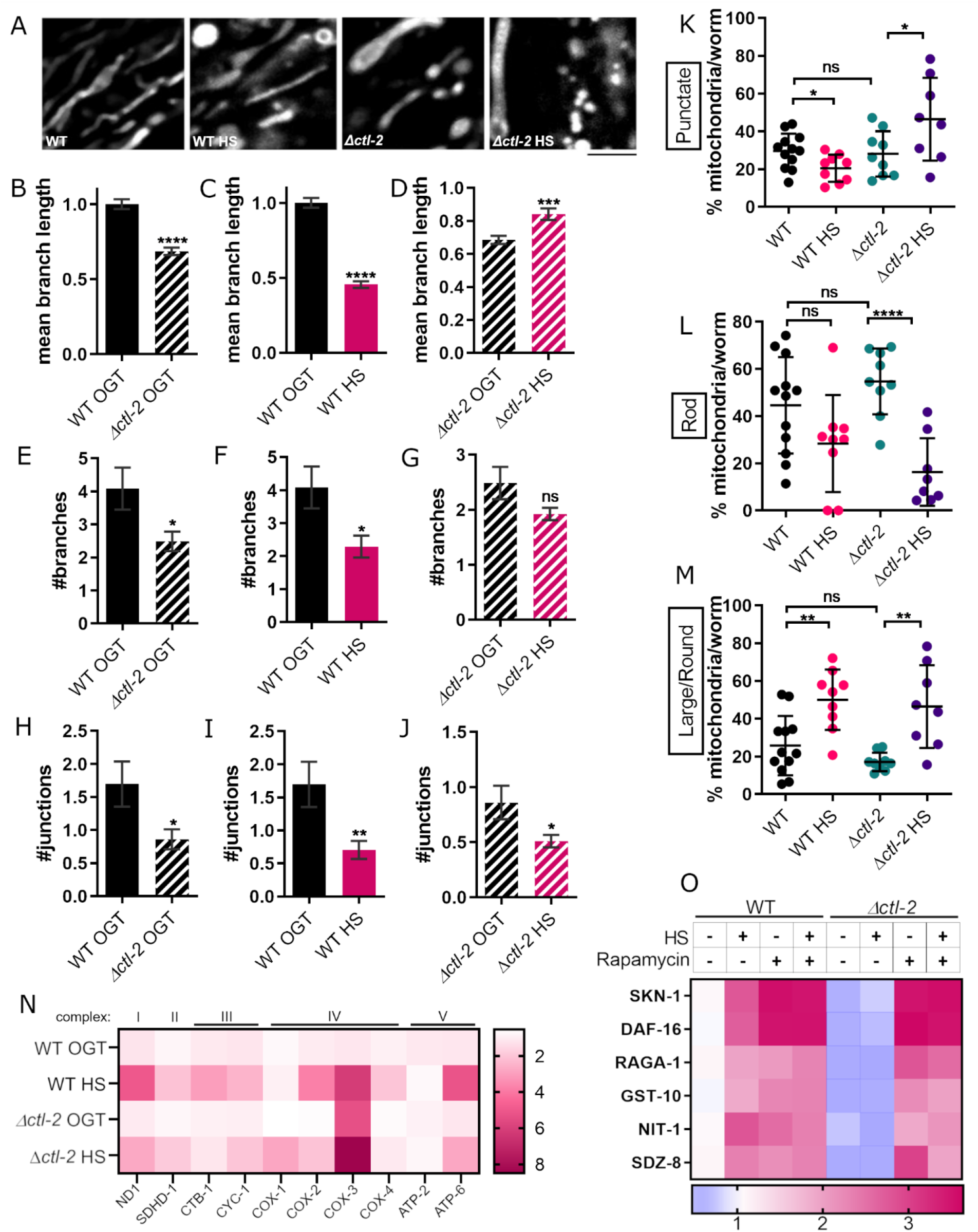
Mitochondrial transcript levels and mitochondrial morphology are altered in the *Δctl-2* mutant. (A) Representative images of MitoTracker stained mitochondria display differences in mitochondrial morphology between WT and *Δctl-2* mutant. (B-J) Analysis and quantification of networked mitochondria reveals that HS results in fewer and shorter branches i.e. less branched mitochondrial networks relative to WT control. *Δctl-2* strain also has less branched mitochondria at OGT. ****P<0.0001; ***P < 0.001; **P < 0.01; *P < 0.05 (t-test). (K-M) Comparison of individual mitochondria in WT and *Δctl-2* strain shows that non-branched mitochondria are similar at OGT in both strains, but during HS, WT displays larger, globular mitochondria, while *Δctl-2* mutant has more punctate mitochondria. (N) Respiratory chain subunits are increased during HS in both strains, but to a lesser extent in the *Δctl-2* strain, except for COX-3 subunit of complex IV. (O) RT-qPCR measurement reveals increased transcript levels of TORC1 downstream targets, indicating TORC1 is inhibited in the WT during HS but not in the peroxisomal mutant. Color of the squares on the heat map corresponds to the mean value of the log fold change. Transcript levels are normalized to ACT-1. ****P<0.0001; ***P < 0.001; **P < 0.01; *P < 0.05 (ANOVA).

Analysis of the non-branched mitochondria reveals further differences (Figure 6K-M). At OGT, long, rod-like mitochondria are most commonly observed. However, at HS, we observed a greater proportion of large and round mitochondria in WT, while *Δctl-2* strain accumulated both large and round mitochondria, as well as small, punctate mitochondria, indicative of active mitochondrial fission. We further measured transcript levels of respiratory chain subunits. During HS, respiratory chain subunit encoding genes were generally upregulated, with a few exceptions; CTB-1 subunit of complex III was not upregulated during HS in *Δctl-2* strain, while complex IV COX-1 (CTC-1) subunit was upregulated only in the *Δctl-2* mutant, approximately 3-fold compared to WT. Furthermore, transcript levels for complex IV subunit COX-3 (CTC-3) were approximately 6 times higher in WT during HS, which was comparable to the level measured in *Δctl-2* strain both at OGT and HS (Figure 6N). While these results suggest the changes in respiratory metabolism during HS, they do not point towards different trends between WT and *Δctl-2* strain.

### TORC1 is inhibited in WT during HS, but not in the Δctl-2 mutant

Finally, we measured TORC1 activity during HS. TORC1 inhibition has been repeatedly demonstrated to extend lifespan across species (Bonawitz, Chatenay-Lapointe et al. 2007, Pan and Shadel 2009, Wei, Fabrizio et al. 2009, Pan, Schroeder et al. 2011, Robida-Stubbs, Glover-Cutter et al. 2012, Johnson, Rabinovitch et al. 2013, Semchyshyn and Valishkevych 2016, Perić, Lovrić et al. 2017, Musa, Perić et al. 2018). When TORC1 is inhibited, its downstream target SKN-1/Nrf is unrepressed and activates transcription of protective genes, such as superoxide dismutases and catalases, as well as TORC1 subunits as a part of a feedback loop. DAF16/FoxO is also activated upon TORC-1 inhibition: both DAF-16/FoxO and SKN-1/Nrf are inhibited by insulin/IGF-like signaling in *C. elegans* (Kenyon 2010, Kenyon 2011, Robida-Stubbs, Glover-Cutter et al. 2012). We measured TORC-1 activity indirectly by RT-qPCR measurement of its downstream targets: SKN-1, DAF-16, RAGA-1, GST-10, NIT-1, SDZ-8. The RT-Qpcr measurement suggested that TORC-1 was inhibited by HS in the WT strain, indicated by increased transcription of TORC-1 downstream targets, including TORC-1 subunit RAGA-1. Treatment with rapamycin, a well-known inhibitor of TORC-1, triggered similar increase in TORC-1 downstream targets, suggesting that, like rapamycin, HS leads to TORC-1 inhibition. Interestingly, HS did not inhibit TORC-1 in the peroxisomal *Δctl-2* mutant, while rapamycin did (Figure 6O). TORC-1 activity in the *Δctl-2* mutant may be contributing to its shortened lifespan and lack of hormetic effect following mild HS.

## Discussion

Hormetic effects following exposure to low dose of stressors have been to date observed in several organisms (Butov, Johnson et al. 2001, Cypser and Johnson 2002, Hercus, Loeschcke et al. 2003, Calabrese, Bachmann et al. 2007, Rattan and Demirovic 2009, Semchyshyn and Valishkevych 2016, Kumsta, Chang et al. 2017). However, the general mechanisms of how these effects arise is still a matter of debate. One suggested mechanism explains that cells upregulate various cellular defense mechanisms such as chaperones in response to the appearance of misfolded proteins during heat stress. This increase in chaperone concentration and improvement of folding conditions is proposed to be the main driving force behind hormetic lifespan extension (Richter, Haslbeck et al. 2010, Morimoto 2011). However, several lines of evidence exist that suggest HSR, while necessary, is not by itself sufficient to support stress induced lifespan extension. In yeast, for example, in order for the mother cell to benefit from heat shock, the diffusion barrier between the mother and the nascent daughter cell needs to be permeated, allowing for diffusion of damaged cell components and aging factors, which would otherwise be retained by the mother, into the daughter cell (Baldi, Bolognesi et al. 2017). Moreover, mitochondrial superoxide is required for stress induced lifespan extension in both yeast and *C. elegans* (Van Raamsdonk and Hekimi 2009, Musa, Perić et al. 2018). Furthermore, budding yeast strains that were not able to switch from fermentative to respiratory growth or increase respiration rate during HS did not display extended lifespan despite upregulation of HSPs (Perić, Lovrić et al. 2017, Musa, Perić et al. 2018). Even HSR itself, which is traditionally discussed in terms of HSF-1, HSP-70, and HSP-90 in a regulatory loop, has been shown to be a part of a more vast and complicated regulatory network that is needed to regulate the HSR in different tissues (Guisbert, Czyz et al. 2013). These observations demonstrate that hormetic effects must be a consequence of proper management and fine-tuning of cellular processes during stress, including canonical stress response pathways, as opposed to increased concentration of chaperones alone.

The results presented here further support this idea. Peroxisomes have not been ascribed any vital roles in the HSR so far. The lack of peroxisomal catalase seems to have impaired the mutant’s ability to respond properly to heat stress, evidenced by lack of lifespan extension, decreased thermotolerance, and a range of metabolic changes when compared with wild type worms, which are not contained to peroxisomes. While we do not know the exact role peroxisomes play in the cascade of cellular events from the onset of HS to lifespan extension, it seems that fully functional peroxisomes are necessary for the hormetic effects to occur. Several possibilities can be speculated about based on the presented results. For example, in the absence of the peroxisomal catalase, it is likely that the H_2_O_2_ accumulates in the peroxisomal matrix and triggers the Fenton reaction when in contact with divalent iron. In this way, peroxisomal machineries may suffer oxidative damage, leading also to their compromised function. Furthermore, changes in the FA metabolism may consequently negatively affect membrane composition, making it more prone to damage by oxidation thus also affecting its fluidity. As peroxisomes are sites of lipid catabolism, providing substrates such as shortened FAs for mitochondria to be used for energy production, disrupting their function may in turn disrupt mitochondrial metabolism as well. For example, the observed decrease in transcript levels of the FA synthesis rate limiting enzyme POD-2 in the catalase deficient mutant could result in a deficit of malonyl-CoA, as well as in the inhibition of transport of FAs to mitochondria (Bowman and Wolfgang 2019). Decreased peroxisomal β-oxidation was also shown to increase mortality in yeast on rich media, as a result of buildup of neutral FAs and TAGs (Titorenko and Terlecky 2011). Furthermore, FA metabolism and lipid droplet dynamics have both been implicated in regulating longevity through modulation of availability of certain FAs and lipophilic hormones such as depfachronic acid and pregnenolone which delay aging in *C. elegans* (Russell and Kahn 2007). While we did not observe buildup of FAs and TAGs in *C. elegans*, we cannot exclude the possibility that some lipid species increased in abundance at the expense of others either following HS or in the absence of the peroxisomal catalase, based on the metabolic changes we observed. However, more precise detection methods would be required to observe these changes.

Finally, the lack of TORC-1 inactivation alongside decreased HSR activation in the peroxisomal mutant highlights the complexity and intersectionality of the cellular stress responses. TORC-1 activity has long been correlated with decreased lifespan across organisms. Together, these data show that stress induced lifespan extension is likely a result of a cell wide coordinated response, alongside robustly activated canonical HSR. Interestingly, while differences between species do exist in how they respond to and benefit from stress, TORC-1 suppression, chaperone upregulation, and tightly regulated, localized increases of ROS seem to be the conserved mechanisms by which eukaryotic cells build up and maintain resistance to stress and prolong their lifespan.

## Conclusion

Hormetic effects of mild, transient heat shock have been described for a range of organisms from yeast to mammals. Our goal was to explore the effects of HS that contribute to the hormetic effect outside of the canonical HSR. Although further experiments are necessary to describe the exact role and function of peroxisomes in the heat-induced hormesis, our results suggest that peroxisomes may have an indispensable contribution. Peroxisomes have a number of vital roles that may affect survival during stressful conditions, including but not limited to their connection with mitochondria, already implicated in many aspects of stress biology. Peroxisomes therefore may hold answers to many yet unanswered questions related to hormesis and stress induced lifespan extension. Finally, better understanding of its role, as well as the more complete picture of heat stress response across the cell and outside of the canonical HSF-1 pathway, is key to applying the principles of lifespan extension we have learned from unicellular eukaryotes to more complex organisms, and eventually, humans. Future research will show how these responses are coordinated and how they contribute to organismal survival and longevity.

## Methods

### Strains

*C. elegans* trains were obtained from Caenorhabditis Genetics Center at the University of Minnesota (https://cgc.umn.edu/). Animals were maintained at 20 °C on NGM media seeded with OP50 *E. coli*. To obtain *Δctl-2* mutant with GFP reporters we crossed GFP-carrying strains (OG497 and VS15) as follows: unsynchronized populations abundant in L4 stage worms of GFP carrying strains were repeatedly heat shocked at 30°C or 37°C for multiple generations until male worms were observed on the plate. The males were then picked and placed on a separate 3 cm plate seeded with OP50 with 2 or 3 L4 hermaphrodites to enrich for males. GFP carrying males were then picked on a separate 0P50 seeded plates together with *Δctl-2* L4 hermaphrodites. Once the F2 worms reached L4 stage, individual F2 worms were picked onto individual plates to lay eggs, after which they were collected for screening under fluorescent microscope. The eggs and the progeny of the worms that exhibited appropriate fluorescence were confirmed to be *Δctl-2* using PCR with the following primers:

(i) ctl-2_Fw TTAGATATGAGAGCGAGCCTGTTTC;

(ii) ctl-2_Rc CTAGTGGTACATCCATGCAAATGC.

### Synchronization

Mixed plates of worms were chunked onto fresh NGM plates seeded with OP50 and grown until they contained predominantly gravid adults. Gravid adults were washed from the plate with M9 buffer or water, and the plate was gently scraped with a spatula to collect the already laid eggs. The eggs and gravid worms were settled by gravity or brief centrifugation at low RPM in 15 mL Falcon tubes. The worm and egg pellet was incubated in 20% hypochlorite solution for 5 minutes with occasional vortexing, until adult hermaphrodites appeared broken. The eggs were then settled by centrifugation and washed with sterile M9 buffer two times. The pellet was then either incubated in a sterile 7 cm plate in 7 mL of sterile M9 buffer at RT with light shaking or seeded on a clean NGM plate where they could hatch without food. After hatching, the arrested L1 larvae were transferred to NGM plates seeded with appropriate food source and incubated at 20°C.

### Lifespan measurement

10-15 L4 hermaphrodites from a synchronized population were transferred to 3 cm NGM plates seeded with OP50. Worms were counted and transferred to a fresh plate every two days until they stopped moving or responding to prodding with the pick. Minimum of 100 worms was picked for each condition for every repetition to ensure sufficient number after censoring stragglers and bagged worms.

### Heat shock

Heat shock was performed on plates at 30°C. Worms were incubated at 20°C until L4 stage, when they were transferred to a 30°C incubator for HS. Following HS, they were either immediately collected for RNA isolation or staining without recovery, or returned to 20°C for lifespan analysis.

### Thermotolerance assay

Plates with synchronized and well fed L4 worms were taken from 20°C and incubated at 37°C for set amount of time, after which they were left to recover at RT overnight. Survival was scored the next day; worms that were not moving and did not respond to stimuli were scored as dead. Minimum of 100 worms were counted for each plate.

### RNA isolation

To isolate RNA, 250 μL of Trizol was added to the worm pellet, vortexed for 30 seconds and incubated at 4°C with shaking for 1 hour. Then 50 μL of chloroform was added, followed by a 30 second centrifugation and 3 minutes at RT. The tubes were then centrifuged for 15 minutes at 10000 rcf at 4°C. The top layer (app. 125 μL) was transferred to a fresh tube and the chloroform addition and centrifugation steps were repeated. 125 μL of 2-propanol was added, the tubes were mixed by inverting to avoid shearing the RNA and incubated at RT for 3 minutes. The tubes were centrifuged at 10000 rcf for 10 minutes and supernatant was carefully decanted without disturbing the RNA pellet. The pellet was washed with 250 μL of 70% RNAse-free ethanol and centrifuged at 10000 rcf for 5 minutes at 4°C. Supernatant was removed and the tubes air-dried inside the hood. Pellets were dissolved in 10 μL RNAse-free water and heated at 65°C for 10 minutes. Concentration was determined using Nanodrop (Thermo Scientific) and the quality was checked on an agarose gel. Samples were stored and kept at -80°C.

### Quantitative real-time PCR

cDNA was synthesized from 1000ng of total RNA using iScript™ cDNA Synthesis Kit (Biorad). The resulting cDNA was diluted 100×, mixed with primer pairs for each gene and SYBRgreen (BioRad). All primer pairs were designed using Primer3 with spliced sequences retrieved from Wormbase (wormbase.org) and were made to span at least two exons. The qPCR reaction was run on a LightCycler 480 (Roche) using 40 cycles, after which the melting curves for each well were determined. Final fold change values were estimated relative to the *act-1* gene in the WT control. List of primers is in Supplementary File 1.

### Live imaging

Synchronized worms were washed off the plates and washed from bacteria with M9 or sterile water. Slides with 2% agarose in M9 pads were prepared fresh before each imaging. Washed worms were immobilized using imidazole, placed on the agarose pad, and covered with a cover slip sealed with nail polish. Imaging was done with temperature-controlled Nikon Ti-E Eclipse inverted/UltraVIEW VoX (Perkin Elmer) spinning disc confocal setup, driven by Volocity software (version 6.3; Perkin Elmer). Unless otherwise specified, images were analysed using ImageJ.

### Mitochondrial morphology

Worms were grown on NGM plates containing 2 μM Mitotracker Deep Red until L4 stage. After washing, worms were immobilized with imidazole and imaged on a temperature-controlled Nikon Ti-E Eclipse inverted/UltraVIEW VoX (Perkin Elmer) spinning disc confocal setup, driven by Volocity software (version 6.3; Perkin Elmer). Mitochondrial morphology was analysed using ImageJ as described in (Valente, Maddalena et al. 2017). In short, images were thresholded and skeletonized for the network analysis. Individual mitochondria were analysed manually from raw images with removed background. Statistical difference between groups was determined using t-test.

### Nile red staining

Nile red stock solution was prepared as a 5 mg/mL solution in acetone. The stock solution was stirred in dark for 2 hours before use. Working solution was made by diluting 6 μL of the stock solution in 1 mL of 40% isopropanol. Worms were washed from the plates and bacteria were removed by washing in PBST. Pellet of washed worms was incubated for 3 minutes at RT with gentle rocking in 40% isopropanol. After incubation worms were pelleted and the supernatant removed. Worm pellet was incubated in 600 μL of working solution for 2 hours in the dark. After incubation, Nile red solution was removed and the worm pellet was washed in PBST for 30 minutes. Worms were imaged on a 2% agarose pad on a Nikon Ti-E Eclipse inverted/UltraVIEW VoX (Perkin Elmer) spinning disc confocal setup, driven by Volocity software (version 6.3; Perkin Elmer). Green fluorescence was recorded. Imaging of unstained control under the same conditions gave no signal.

### Oil red O staining

Oil Red O stock solution (ORO) was prepared as a 5mg/mL solution in 100% isopropanol. Working solution was made by diluting stock 3:2 in water (60% isopropanol final concentration). The working solution was filtered through a 0.2 μm filter and allowed to mix for 2 hours before use. Worms were washed from the plates and bacteria were removed by washing in PBST. Pellet of washed worms was incubated for 3 minutes at RT with gentle rocking in 40% isopropanol. After incubation, worms were pelleted and the supernatant was removed. The worm pellet was incubated for 2 hours in 600 μL of ORO working solution with gentle mixing. After staining, the pellet was washed for 30 minutes in PBST. Imaging was done with a color capable camera.

### ROS measurement

Worms were washed from the bacteria until the supernatant was clear. Washed worms were fixed in 3.7% formaldehyde for 15 minutes. Fixed worms were pelleted and washed in M9. The worm pellet was stained with 5 μM Cell ROX Deep Red for 1 hour. Stained worms were imaged within 2 hours and red fluorescence proportional to amount of ROS was quantified with ImageJ. Statistical difference was determined using multiple t-tests.

### Statistical analysis

Unless otherwise stated, data in graphs are mean ± SEM from three biological and three technical replicates. Survival experiments were tested with log-rank (Mantel-Cox) test. ANOVA and t-test were used for all other measurements, as indicated for individual graphs. Graphing and statistical analysis was done using GraphPad Prism and Minitab software. ****P <0.0001; ***P < 0.001; **P < 0.01; *P < 0.05.

## Acknowledgements

A.K. is supported by the Heisenberg grant from the Deutsche Forschungsgemeinschaft. M.M. was supported by the Mediterranean Institute for Life Sciences, as well as by FEBS and EMBO short-term fellowships. Grant 337327 from the European Research Council to N.R.; I.M. is supported by the John Black Charitable Foundation and the John Fell Funds.

The authors would like to thank Tea Copić, Dirk Schwitters and Marina Konta for their excellent technical contribution.

## Notes

### Competing Interest Statement

The authors have declared no competing interest.

## Bibliography

Baldi, S., A. Bolognesi, A. C. Meinema and Y. Barral (2017). “Heat stress promotes longevity in budding yeast by relaxing the confinement of age-promoting factors in the mother cell.” Elife 6.

Ben-Zvi, A., E. A. Miller and R. I. Morimoto (2009). “Collapse of proteostasis represents an early molecular event in Caenorhabditis elegans aging.” Proc Natl Acad Sci U S A 106(35): 14914–14919.

Bonawitz, N. D., M. Chatenay-Lapointe, Y. Pan and G. S. Shadel (2007). “Reduced TOR signaling extends chronological life span via increased respiration and upregulation of mitochondrial gene expression.” Cell Metab 5(4): 265–277.

Bowman, C. E. and M. J. Wolfgang (2019). “Role of the malonyl-CoA synthetase ACSF3 in mitochondrial metabolism.” Adv Biol Regul 71: 34–40.

Butov, A., T. Johnson, J. Cypser, I. Sannikov, M. Volkov, M. Sehl and A. Yashin (2001). “Hormesis and debilitation effects in stress experiments using the nematode worm Caenorhabditis elegans: the model of balance between cell damage and HSP levels.” Exp Gerontol 37(1): 57–66.

Calabrese, E. J., K. A. Bachmann, A. J. Bailer, P. M. Bolger, J. Borak, L. Cai, N. Cedergreen, M. G. Cherian, C. C. Chiueh, T. W. Clarkson, R. R. Cook, D. M. Diamond, D. J. Doolittle, M. A. Dorato, S. O. Duke, L. Feinendegen, D. E. Gardner, R. W. Hart, K. L. Hastings, A. W. Hayes, G. R. Hoffmann, J. A. Ives, Z. Jaworowski, T. E. Johnson, W. B. Jonas, N. E. Kaminski, J. G. Keller, J. E. Klaunig, T. B. Knudsen, W. J. Kozumbo, T. Lettieri, S. Z. Liu, A. Maisseu, K. I. Maynard, E. J. Masoro, R. O. McClellan, H. M. Mehendale, C. Mothersill, D. B. Newlin, H. N. Nigg, F. W. Oehme, R. F. Phalen, M. A. Philbert, S. I. Rattan, J. E. Riviere, J. Rodricks, R. M. Sapolsky, B. R. Scott, C. Seymour, D. A. Sinclair, J. Smith-Sonneborn, E. T. Snow, L. Spear, D. E. Stevenson, Y. Thomas, M. Tubiana, G. M. Williams and M. P. Mattson (2007). “Biological stress response terminology: Integrating the concepts of adaptive response and preconditioning stress within a hormetic dose-response framework.” Toxicol Appl Pharmacol 222(1): 122–128.

Cox, C. S., S. E. McKay, M. A. Holmbeck, B. E. Christian, A. C. Scortea, A. J. Tsay, L. E. Newman and G. S. Shadel (2018). “Mitohormesis in Mice via Sustained Basal Activation of Mitochondrial and Antioxidant Signaling.” Cell Metab 28(5): 776–786.e775.

Cypser, J. R. and T. E. Johnson (2002). “Multiple stressors in Caenorhabditis elegans induce stress hormesis and extended longevity.” J Gerontol A Biol Sci Med Sci 57(3): B109–114.

Guisbert, E., D. M. Czyz, K. Richter, P. D. McMullen and R. I. Morimoto (2013). “Identification of a tissue-selective heat shock response regulatory network.” PLoS Genet 9(4): e1003466.

Hansen, M., T. Flatt and H. Aguilaniu (2013). “Reproduction, fat metabolism, and life span: what is the connection?” Cell Metab 17(1): 10–19.

Hercus, M. J., V. Loeschcke and S. I. Rattan (2003). “Lifespan extension of Drosophila melanogaster through hormesis by repeated mild heat stress.” Biogerontology 4(3): 149–156.

Johnson, S. C., P. S. Rabinovitch and M. Kaeberlein (2013). “mTOR is a key modulator of ageing and age-related disease.” Nature 493(7432): 338–345.

Kenyon, C. (2011). “The first long-lived mutants: discovery of the insulin/IGF-1 pathway for ageing.” Philos Trans R Soc Lond B Biol Sci 366(1561): 9–16.

Kenyon, C. J. (2010). “The genetics of ageing.” Nature 464(7288): 504–512.

Kim, H. E., A. R. Grant, M. S. Simic, R. A. Kohnz, D. K. Nomura, J. Durieux, C. E. Riera, M. Sanchez, E. Kapernick, S. Wolff and A. Dillin (2016). “Lipid Biosynthesis Coordinates a Mitochondrial-to-Cytosolic Stress Response.” Cell 166(6): 1539–1552.e1516.

Kumsta, C., J. T. Chang, J. Schmalz and M. Hansen (2017). “Hormetic heat stress and HSF-1 induce autophagy to improve survival and proteostasis in C. elegans.” Nat Commun 8: 14337.

Lapierre, L. R., S. Gelino, A. Meléndez and M. Hansen (2011). “Autophagy and lipid metabolism coordinately modulate life span in germline-less C. elegans.” Curr Biol 21(18): 1507–1514.

López-Martínez, G. and D. A. Hahn (2014). “Early life hormetic treatments decrease irradiation-induced oxidative damage, increase longevity, and enhance sexual performance during old age in the Caribbean fruit fly.” PLoS One 9(1): e88128.

Mendillo, M. L., S. Santagata, M. Koeva, G. W. Bell, R. Hu, R. M. Tamimi, E. Fraenkel, T. A. Ince, L. Whitesell and S. Lindquist (2012). “HSF1 drives a transcriptional program distinct from heat shock to support highly malignant human cancers.” Cell 150(3): 549–562.

Morimoto, R. I. (2008). “Proteotoxic stress and inducible chaperone networks in neurodegenerative disease and aging.” Genes Dev 22(11): 1427–1438.

Morimoto, R. I. (2011). “The heat shock response: systems biology of proteotoxic stress in aging and disease.” Cold Spring Harb Symp Quant Biol 76: 91–99.

Morton, E. A. and T. Lamitina (2013). “Caenorhabditis elegans HSF-1 is an essential nuclear protein that forms stress granule-like structures following heat shock.” Aging Cell 12(1): 112–120.

Musa, M., M. Perić, P. Bou Dib, S. Sobočanec, A. Šarić, A. Lovrić, M. Rudan, A. Nikolić, I. Milosević, K. Vlahoviček, N. Raimundo and A. Kriško (2018). “Heat-induced longevity in budding yeast requires respiratory metabolism and glutathione recycling.” Aging (Albany NY) 10(9): 2407–2427.

Olsen, A., M. C. Vantipalli and G. J. Lithgow (2006). “Lifespan extension of Caenorhabditis elegans following repeated mild hormetic heat treatments.” Biogerontology 7(4): 221–230.

Pan, Y., E. A. Schroeder, A. Ocampo, A. Barrientos and G. S. Shadel (2011). “Regulation of yeast chronological life span by TORC1 via adaptive mitochondrial ROS signaling.” Cell Metab 13(6): 668–678.

Pan, Y. and G. S. Shadel (2009). “Extension of chronological life span by reduced TOR signaling requires down-regulation of Sch9p and involves increased mitochondrial OXPHOS complex density.” Aging (Albany NY) 1(1): 131–145.

Perić, M., A. Lovrić, A. Šarić, M. Musa, P. Bou Dib, M. Rudan, A. Nikolić, S. Sobočanec, A. M. Mikecin, S. Dennerlein, I. Milošević, K. Vlahoviček, N. Raimundo and A. Kriško (2017). “TORC1-mediated sensing of chaperone activity alters glucose metabolism and extends lifespan.” Aging Cell 16(5): 994–1005.

Petriv, O. I. and R. A. Rachubinski (2004). “Lack of peroxisomal catalase causes a progeric phenotype in Caenorhabditis elegans.” J Biol Chem 279(19): 19996–20001.

Prahlad, V., T. Cornelius and R. I. Morimoto (2008). “Regulation of the cellular heat shock response in Caenorhabditis elegans by thermosensory neurons.” Science 320(5877): 811–814.

Rattan, S. I. and D. Demirovic (2009). “Hormesis can and does work in humans.” Dose Response 8(1): 58–63.

Rea, S. L., D. Wu, J. R. Cypser, J. W. Vaupel and T. E. Johnson (2005). “A stress-sensitive reporter predicts longevity in isogenic populations of Caenorhabditis elegans.” Nat Genet 37(8): 894–898.

Richter, K., M. Haslbeck and J. Buchner (2010). “The heat shock response: life on the verge of death.” Mol Cell 40(2): 253–266.

Robida-Stubbs, S., K. Glover-Cutter, D. W. Lamming, M. Mizunuma, S. D. Narasimhan, E. Neumann-Haefelin, D. M. Sabatini and T. K. Blackwell (2012). “TOR signaling and rapamycin influence longevity by regulating SKN-1/Nrf and DAF-16/FoxO.” Cell Metab 15(5): 713–724.

Russell, S. J. and C. R. Kahn (2007). “Endocrine regulation of ageing.” Nat Rev Mol Cell Biol 8(9): 681–691.

Scott, B. R. (2014). “Radiation-hormesis phenotypes, the related mechanisms and implications for disease prevention and therapy.” J Cell Commun Signal 8(4): 341–352.

Semchyshyn, H. M. and B. V. Valishkevych (2016). “Hormetic Effect of H2O2 in Saccharomyces cerevisiae: Involvement of TOR and Glutathione Reductase.” Dose Response 14(2): 1559325816636130.

Shama, S., C. Y. Lai, J. M. Antoniazzi, J. C. Jiang and S. M. Jazwinski (1998). “Heat stress-induced life span extension in yeast.” Exp Cell Res 245(2): 379–388.

Stincone, A., A. Prigione, T. Cramer, M. M. Wamelink, K. Campbell, E. Cheung, V. Olin-Sandoval, N. M. Grüning, A. Krüger, M. Tauqeer Alam, M. A. Keller, M. Breitenbach, K. M. Brindle, J. D. Rabinowitz and M. Ralser (2015). “The return of metabolism: biochemistry and physiology of the pentose phosphate pathway.” Biol Rev Camb Philos Soc 90(3): 927–963.

Takeuchi, T., M. Suzuki, N. Fujikake, H. A. Popiel, H. Kikuchi, S. Futaki, K. Wada and Y. Nagai (2015). “Intercellular chaperone transmission via exosomes contributes to maintenance of protein homeostasis at the organismal level.” Proc Natl Acad Sci U S A 112(19): E2497–2506.

Titorenko, V. I. and S. R. Terlecky (2011). “Peroxisome metabolism and cellular aging.” Traffic 12(3): 252–259.

Valente, A. J., L. A. Maddalena, E. L. Robb, F. Moradi and J. A. Stuart (2017). “A simple ImageJ macro tool for analyzing mitochondrial network morphology in mammalian cell culture.” Acta Histochem 119(3): 315–326.

van Oosten-Hawle, P. and R. I. Morimoto (2014). “Organismal proteostasis: role of cell-nonautonomous regulation and transcellular chaperone signaling.” Genes Dev 28(14): 1533–1543.

van Oosten-Hawle, P., R. S. Porter and R. I. Morimoto (2013). “Regulation of organismal proteostasis by transcellular chaperone signaling.” Cell 153(6): 1366–1378.

Van Raamsdonk, J. M. and S. Hekimi (2009). “Deletion of the mitochondrial superoxide dismutase sod-2 extends lifespan in Caenorhabditis elegans.” PLoS Genet 5(2): e1000361.

Wei, M., P. Fabrizio, F. Madia, J. Hu, H. Ge, L. M. Li and V. D. Longo (2009). “Tor1/Sch9-regulated carbon source substitution is as effective as calorie restriction in life span extension.” PLoS Genet 5(5): e1000467.

Yang, W. and S. Hekimi (2010). “A mitochondrial superoxide signal triggers increased longevity in Caenorhabditis elegans.” PLoS Biol 8(12): e1000556.

