## Supplemental Table 1 for "Functional peroxisomes are required for heat shock-induced hormesis in *Caenorhabditis elegans*"

Table S1.

| | Fw | Rc |
|---|---|---|
| act-1 | AGGAGTCATGGTCGGTATGG | GCTTCAGTGAGGAGGACTGG |
| nit-1 | CACGAACTCCAGAAGGAAGG | TCGGTGCTTTCCAAGGTAAC |
| raga-1 | AGCTATTTTGGATGCCGATG | TGAATGCGGAGAATTGACTG |
| skn-1 | CTCCATTCGGTAGAGGACCA | GGCGCTACTGTCGATTTCTC |
| daf-16 | AATTGCTCCACCACCATCAT | AATTCCAGGCAGTGGAGATG |
| gst-10 | ATTCGAAGACATTCGGTTCG | AACATGTCGAGGAAGGTTGC |
| sdz-8 | GAGCTACGGAACGAATACGG | CAGTGATGGAACCAAGTCCA |
| pod-2 | CTTGAATGGTCATGGCACAG | ATAGTCGCGGAGCAGTTGAT |
| fasn-1 | TCCATTTGCAACTGATTCCA | ATCCAATTTGATTGGCTTGG |
| fat-5 | GAGGATCCGGTGCTGATG | AGCAGAAGATTCCGACCAAG |
| fat-6 | CTCGTTCAAACATCGCAAAA | GAGCTGGTAGAGTCCGATGG |
| Pdi-6 | GAACCTTTGGGGATGGATTT | GAAGTGCGTCCCTTTCCATA |
| hsp-60 | GTCGGAACCACTGGAGTCAT | CCGGCACAATATCCTGAACT |
| clpp-1 | GGTGAAAAAGGAATGCGAAG | TCACGGTCAAGGGTTTTCTC |
| hsp-6 | GAGCTCCAGCCAAAAGACAG | AATGCTCCAACCTGAGATGG |
| ND1 | CCATCCGTGCTAGAAGACAA | TCCCTTTCACCTTCAGAAAAA |
| sdhd-1 | TGGAATGCTCCCAATTCTTC | CGAGCAGAAGACCAGTGATG |
| CYTB | AATGGGGCCAGGTTATTTTT | TGGCCCTCAAATTGGAATAA |
| cyc-1 | TGGAGTGAAGGTTGATGACG | GAGCCCACTTCTTTCTGGTG |
| cox-1 | TCGGTGGTTTTGGTAACTGA | TGCCCCATTGTTCTTAAAGG |
| cox-2 | TGTTATTCATGCTTGGGCATT | TGCTCCACAAATCTCTGAACAT |
| cox-3 | TTGGCAGCTTATTTTACAGGAA | TCCAACCCCAGATGATGATT |
| cox-4 | GGATACGGAGCAAATGGAGA | GAATTCGGACAGCGTTTGAC |
| atp-2 | GAACCTTCCACCAATCCTCA | TACGGCCAAGAGTTTCTGGT |
| atp-6 | TGTCCTTGTGGAATGGTTGA | CGCACTGTTAAAGCAAGTGG |
| sod-1 | ATCCGAGATCCGTCACGTAG | GCGTTTCCAGTCTTCTTGGA |
| sod-2 | TTGTTCAACCGATCACAGGA | TGACGTTTCCTTTGGAGACC |
| sod-3 | CTTGGCTAAGGATGGTGGAG | TTCCAAAGGATCCTGGTTTG |
| sod-5 | GGAACTGCTGTCTTCGGAAC | CGGAATCTCTTCCTCCATGA |
| prx-5 | GCAAACCCTTTCACAACCAT | AACATTTTTGGAACGCTTGC |
| prx-11 | GGAATTCCTCACAGGAGCAG | TGCGATTTTTCGATTCCTTT |
| gspd-1 | CCAGCAAGTTTGAATGCTGA | CAGCAAGAGCATACGTTGGA |
| daf-22 | AGGTGATCAATGCCCGTAAG | TTGAATTCTCAGCGAACACG |
| maoc-1 | GGGCTGGAAATGATTCTGAC | ACTGTTGGAAGTTCGGAAGC |
| acox-1 | AGCGATCTCCTCCATCTTCA | CGGCCTAGGACAGATCTCAG |
| hsp-16.1 | CGCAGTTCAAGCCAGAAGAT | AATCGCTTCCTTCTTTGGTG |
| hsp-16.2 | CAGCTCAACGTTCCGTTTTT | GATAGCGTACGACCATCCAAA |
| hsp-70 | GCTGATCTTTTCCGCAAGAC | CCAAAGGCTACTGCTTCGTC |
| crt-1 | TTTGTGGCAGGTCAAGTCAG | CTTCTTGGCCTCCTTCTCCT |
| uggt-1 | CGGATATGTGCCATTCTGTG | TTTGGATCGCCAGATAGTCC |
